# Monomerization of Far-Red Fluorescent Proteins

**DOI:** 10.1101/162842

**Authors:** Timothy M. Wannier, Sarah Gillespie, Nicholas Hutchins, R. Scott McIsaac, Kevin S. Brown, Stephen L. Mayo

## Abstract

*Anthozoa* class red fluorescent proteins (RFPs) are frequently used as biological markers, with far-red emitting variants (λ_em_ ~ 600 – 900 nm) sought for whole animal imaging because biological tissues are permeable to light in this range. A barrier to the use of naturally occurring RFP variants as molecular markers is that all are tetrameric, which is not ideal for cell biological applications. Efforts to engineer monomeric RFPs have usually produced dimmer and blue-shifted variants, as the chromophore is sensitive to small structural perturbations. In fact, despite much effort, only four native RFPs have been successfully monomerized, leaving the vast majority of RFP biodiversity untapped in biomarker development. Here we report the generation of monomeric variants of HcRed and mCardinal, both far-red dimers, and describe a comprehensive methodology for the rapid monomerization of novel red-shifted oligomeric RFPs. Among the resultant variants, is mKelly1 (emission maximum: λ_em_ = 656 nm), which along with the recently reported mGarnet2 (1), forms a new class of bright, monomeric, far-red FPs.

## Introduction

The development of red fluorescent proteins (RFPs) as tags for molecular imaging has long focused on monomerization, increased brightness, and pushing excitation and emission to ever-longer wavelengths. These traits are desirable for live animal imaging, as far-red to near infrared light penetrates tissue with minimal absorption in what is known as the near infrared window (~625–1300 nm) (2, 3). Monomericity is important because oligomerization of an FP tag can artificially aggregate its linked protein target, altering diffusion rates and interfering with target transport, trafficking, and activity (4, 5). Recently a new class of infrared fluorescent proteins (iRFPs) was developed from the bacterial phytochrome (6), but these require the covalent linkage of a small molecule chromophore, biliverdin, limiting their use to cells and organisms that make this molecule in sufficient quantity. *Anthozoa* class RFPs (such as mCherry and mKate) have the advantage that the chromophore is created via a self-processing reaction, necessitating only molecular O_2_ for chromophore formation.

To our knowledge, ~50 native RFPs and ~40 chromoproteins (CPs) with peak absorbance in the red or far-red (absorbance maximum: λ_abs_ >550 nm) have been described to date, but most have not been extensively characterized because they are as a class tetrameric, and thus are less useful as biological markers (7, 8). An underlying biological reason for the obligate tetramerization of native RFPs has been hinted at, but is not well understood (9-12). Oligomerization does seem to play an important structural role, however, as breaking tetramerization without abrogating fluorescence has proved difficult, and successful monomerization has always led to either a hypsochromic shift to λ_em_ or a decrease in brightness (13-16). Previous efforts to monomerize native RFP tetramers have relied on lengthy engineering trajectories, and have been successful in only four cases (Table 1). Generally, mutations are first introduced into tight interfaces to weaken oligomerization, an inefficient process that impairs fluorescence, and then random mutagenesis and screening isolate partially recovered variants. After many such cycles monomeric variants have been found, but protein core and chromophore-proximal mutations are invariably introduced, making it difficult to exert any significant degree of control over the fluorescent properties of the resultant monomer. It is thus difficult to know whether the poor spectroscopic characteristics of engineered monomers are an unavoidable consequence of monomerization or only the manifestation of a suboptimal evolutionary path.

**Table 1.** Previously monomerized first generation RFPs

| Monomeric RFP | Brightness ($\Phi \times \epsilon$) / 1000 | $\lambda_{em}$ (nm) | Mutations to Core | Total Mutations | Immediate Parent (dimer/tetramer) | Brightness ($\Phi \times \epsilon$) / 1000 | $\lambda_{em}$ (nm) | Mutations to Core | Total Mutations |
| --- | --- | --- | --- | --- | --- | --- | --- | --- | --- |
| mRFP1 | 12.5 | 607 | 13 | 33 | DsRed (T) | 59.3 | 583 | n/a | n/a |
| DsRed.M1 | 3.5 | 586 | 10 | 45 | " | " | " | " | " |
| FusionRed | 18.0 | 608 | 9 | 45 | mKate2 (D) | 18.0 | 630 | 7 | 27 |
| mRuby | 39.2 | 605 | 6 | 40 | eqFP611 (T) | 35.1 | 611 | n/a | n/a |
| mKeima | 3.5 | 620 | 7 | 17 | dKeima (D) | 7.6 | 616 | 5 | 13 |
| mGinger1 | 1.2 | 637 | 7 | 45 | HcRed-7 (D) | 4.8 | 643 | 4 | 8 |
| mGinger2 | 1.5 | 631 | 7 | 49 | " | " | " | " | " |
| mKelly1 | 5.7 | 656 | 15 | 52 | mCardinal (D) | 9.6 | 658 | 15 | 44 |
| mKelly2 | 6.5 | 649 | 15 | 52 | " | " | " | " | " |

Here we present a comprehensive engineering strategy for the monomerization of novel RFPs that differentiates itself by treating separately the problems of protein stabilization, core optimization, and surface design. We sample mutational space both stochastically, through error-prone mutagenesis, and rationally, by analysis of multiple sequence alignments (MSAs) and computational protein design (CPD). Two far-red oligomeric proteins were targeted for monomerization: HcRed (λ_em_ = 633 nm), a dimer/tetramer (17), and mCardinal (λ_em_ = 658 nm), a reported monomer that we have confirmed to in fact be dimeric. The monomeric RFPs reported here include two monomeric HcRed variants: mGinger1 (λ_em_ = 637 nm) and mGinger2 (λ_em_ = 631), and two monomeric mCardinal variants: mKelly1 (λ_em_ = 648 nm) and mKelly2 (λ_em_ = 643 nm), which are among the brightest far-red monomeric FPs to have been reported.

## Results

### Step-wise monomerization of HcRed

We first chose HcRed, a far-red FP that has been engineered but never successfully monomerized (17, 18). As we have previously demonstrated that oligomericity and brightness can be treated as separate protein design problems (19), we devised a workflow that separately targets the chromophore environment—to engineer a protein core that maintains structural integrity absent stabilizing oligomeric interactions—and the protein surface—to drive monomerization. *Anthozoa* class RFPs have two oligomeric interfaces, named AB and AC (20), with the AC interface being the more stable of the two and burying a large hydrophobic surface (21). Early engineering to HcRed partially disrupted oligomerization at the AB interface, but all mutations to the AC interface were found to vitiate fluorescence. To test the integrity of the AC interface, we made successive deletions to HcRed’s C-terminal tail (residues 219-227), which plays an integral role in the AC interaction (Figure 1A). HcRed lost significant brightness with the deletion of just one C-terminal residue, and was non-fluorescent after any further deletion, demonstrating that optimization would be necessary prior to monomerization.

**Figure 1.**
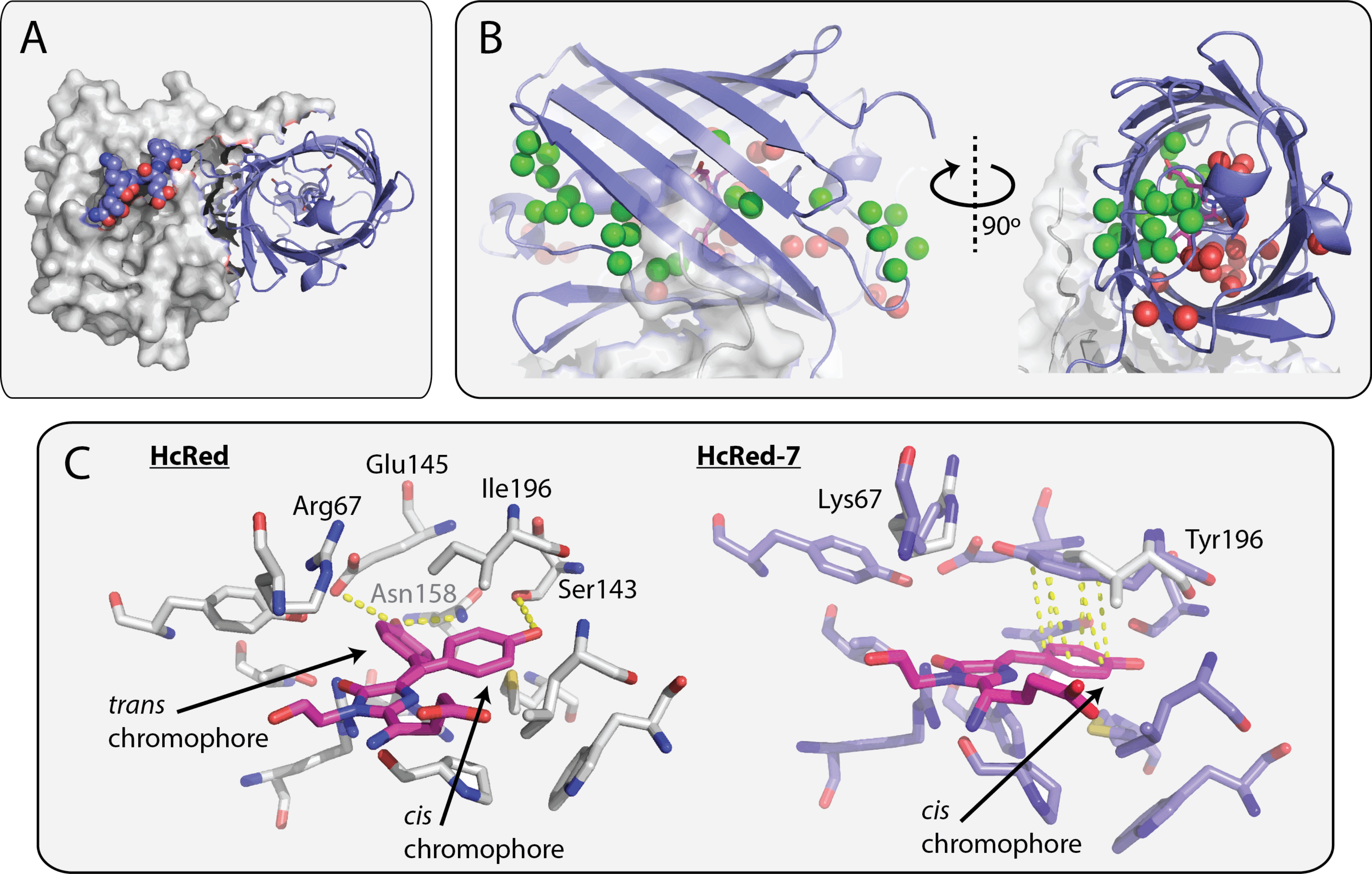
The structure of HcRed7, a far-red dimer. (*A*) The HcRed7 dimeric interface is stabilized by its C-terminal tail. One monomer is shown as a cartoon while the second is shown as a surface; residues 222-227 are shown in spheres. (*B*) A water channel stretches from a structural flaw in HcRed7’s β-barrel through the center of the protein to a smaller defect at its outlet on the other end of the barrel. The C-terminal tail/AC interface inter-molecular interaction bisects this channel, potentially stabilizing it. (C) The crystal structure of HcRed (PDB-ID: 1YZW – in gray sticks) shows dual occupancy of the chromophore’s phenolate group. The cis chromophore is stabilized by a C143S mutation from parent protein HcCP. The trans chromophore is stabilized by two hydrogen bonds (in yellow) from Glu145 and Asn158. Two mutations (R67K and I196Y) were made to HcRed to create HcRed7 (slate sticks). Tyr196 in HcRed7 stabilizes the *cis* chromophore with a π-stacking interaction (in yellow). Lys67 is an important catalytic residue.

So, we endeavored to engineer a more stable core, identifying two mutational hotspots from an alignment of far-red RFPs: (A) a group of residues that surrounds alternative conformations of the chromophore’s phenolate ring and (B) a region above the plane of the chromophore, between the central α-helix and the unbroken AC oligomeric interface (Figure S1). Generally in RFPs, the *cis* chromophore—the phenolate ring sits *cis* to the proximal nitrogen on the imidazolinone ring rather than *trans* to it—is the fluorescent species (22). In engineering HcRed from its chromoprotein parent HcCP, the *cis* chromophore was stabilized over the non-fluorescent *trans* chromophore by way of a cysteine to serine mutation at position 143, which provides a hydrogen bond to the cis phenolate oxygen (Figure 1C). We reasoned that further stabilization of the *cis* chromophore would increase brightness, and so designed a first core library (cLibA) to target hotspot A, mutating *trans*-stabilizing amino acids, placing bulkier side chains into the *trans* pocket, and allowing varied hydrogen bonding geometries to the *cis* chromophore. A second core library (cLibB) targeted hotspot B along with two chromophore-backing positions (Gly28 and Met41 are implicated in maturation and color) (21, 23, 24). Two key features of this hotspot are a channel populated by structural water molecules that stretches to the protein surface, and Arg67, a key catalytic residue. Mutations to this region may serve to occlude access to the chromophore by bulk solvent upon monomerization, and to allow room for chromophore processing. Small libraries of < 1,000 protein variants were guided by the far-red RFP alignment (Table S1), and after screening each library to > 95% coverage on large LB agar plates supplemented with IPTG, we fully characterized 16 cLibA variants and 21 cLibB variants. The variants showed brightness increases of up to ten-fold and displayed an incredible range of emission profiles, with λ_em_ between 606 and 647 nm. To determine which if any variants would be amenable to monomerization, we tested a five-residue tail deletion. Eight variants showed detectable fluorescence after the tail deletion, with a double mutant (HcRed7: R67K/I196Y) being the most red-shifted (λ_em_ = 642 nm). The core mutations in HcRed7 bathochromically shift its emission by 9 nm, improve its quantum yield (Φ) by 60%, and thermostabilize the protein by 6 °C. HcRed7, however, loses significant brightness with the deletion of a sixth tail residue – HcRed7Δ6) and becomes 16°C less thermostable, indicating that the protein is not wholly optimized for monomerization (Table 2).

**Table 2.** Important proteins

| RFP | $\Phi$ | $\epsilon (M^{-1} cm^{-1}) / 1000$ | Brightness<br>($\Phi \times \epsilon$) /<br>1000 | $\lambda_{ex}$<br>(nm) | $\lambda_{em}$<br>(nm) | Apparent<br>$T_m$ (°C) |
| --- | --- | --- | --- | --- | --- | --- |
| HcRed | 0.05 | 70 | 6.0 | 585 | 633 | 69.0 |
| HcRed-7 | 0.08 | 75 | 8.4 | 592 | 645 | 75.0 |
| HcRed-7 $\Delta 5$ | 0.06 | 69 | 4.3 | 592 | 643 | 70.5 |
| HcRed-7 $\Delta 6$ | | | | 582 | 635 | 65.0 |
| HcRed-77 | 0.05 | † | † |  |  | 67.5 |
| HcRed-m1 | 0.01 | † | † |  |  | 64.0 |
| HcRed-m114 | 0.02 | † | † |  |  | 58.0 |
| mGinger1 | 0.02 | 58 | 1.2 | 587 | 637 | 79.0 |
| mGinger2 | 0.04 | 36 | 1.5 | 578 | 631 | 80.0 |
| mCardinal | 0.12 | 80 | 9.6 | 601 | 658 |  |
| mCardinal$\Delta 20$ | | | | | | |
| mCardinal-mut6 | 0.13 | 60 | 7.8 |  |  |  |
| mKelly1 | 0.13 | 44 | 5.7 | 596 | 656 |  |
| mKelly2 | 0.15 | 43 | 6.5 | 598 | 649 |  |

† - Extinction coefficient (and therefore brightness) could not be measured because of multiple chromophore species present.

To further optimize HcRed7Δ6 for monomerization, we took aim at improving the thermo-stability of the protein Thermostability has been shown to increase a protein’s evolvability (25) and consensus design is one of the best tools for improving thermostability (26). We constructed a large MSA that consists of every *Aequorea victoria* class FP; a total of 741 sequences (see supplemental Methods), and then built a library to sample all 105 non-consensus positions in HcRed with the consensus amino acid, and compared this to a strategy of error-prone mutagenesis. We screened the consensus (~1.2 mutations per variant) and error-prone (~1.8 mutations per variant) libraries at 675 nm to allow maximal differentiation between far-red variants whose λ_em_ was between 630-640 nm and a large population of near-red variants whose emission peaked between 605-620 nm, but which were often brighter. The consensus library was screened to 40x coverage (~4,300 clones) and ~8,600 clones were screened from the error-prone library. Consensus library variants significantly outperformed error-prone library variants (Figure S2), and so we combined seven of the top consensus variants together into a chimeric protein, HcRed77, which recovered all of HcRed7Δ6’s lost brightness and much of its thermostability.

Finally, to monomerize HcRed77 we targeted the AC interface with a CPD procedure that we describe in previous work (19). We focused on a set of five hydrophobic residues (Val146, Val159, Ile170, Phe191, and Phe193) at the heart of the AC interface that make extensive intermolecular contacts (Figure S3), and built a small combinatorial library guided by the design. We isolated a first-generation monomer: HcRedm1 and verified it to be monomeric by fast protein liquid chromatography (FPLC) and analytical ultracentrifugation (AUC) (Figure 2), but the protein was dim and expressed poorly. We attributed these poor attributes to incomplete thermo-stabilization of HcRed77, and so we screened mutations from the error-prone library via DNA shuffling and then increased the temperature of screening from 30°C to 37°C for a final round of error-prone mutagenesis and isolated two bright variants with improved brightness and thermostability higher than the parent HcRed7: mGinger1 and mGinger2 (Table 2).

**Figure 2.**
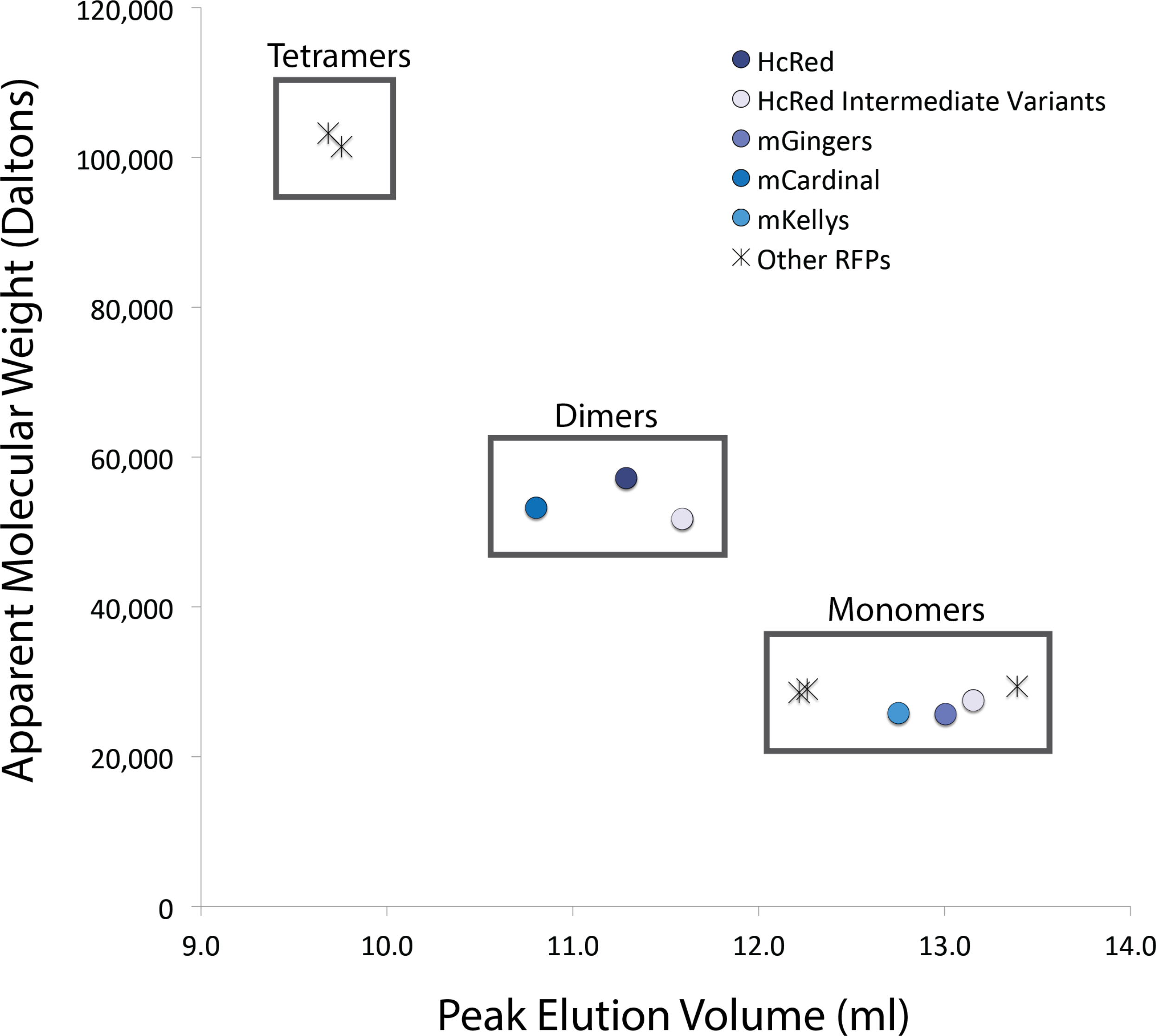
Oligomeric analysis of RFPs. The apparent molecular weight as calculated from a c(M) distribution of sedimentation velocity data from an analytical ultracentrifuge run is plotted on the Y-axis. The X-axis shows the peak elution volume as measured at 590 nm absorbance by size exclusion chromatography. Clear groupings are boxed as monomers, dimers, and tetramers.

### Two-step monomerization of mCardinal

The monomerization of HcRed required three design elements: core optimization, protein thermo-stabilization, and surface design; and pulled mutational diversity from three sources: a large MSA, CPD, and error-prone mutagenesis. While the engineering process for the development of the mGingers was rational and involved a shorter trajectory than past procedures, we felt that it could be further improved by integrating the three design objectives into one large library. We targeted mCardinal, a recently reported variant of mNeptune that was reported to be monomeric, but which we have shown to be dimeric by both FPLC and AUC (Figure 2). In fact, the crystal structure of mCardinal (4OQW) shows the protein adopting a classic tetrameric RFP conformation, similar to DsRed and mCardinal’s wild-type progenitor, eqFP578 (27).

First, as with HcRed, we probed tail deletion variants of mCardinal, which was engineered to have a long, 20-AA C-terminal tail. The first 15 residues were easily removed (equivalent to HcRedΔ4), but as with HcRed, mCardinalΔ16 is significantly dimmer and mCardinalΔ18 is essentially non-fluorescent. To isolate error-prone mutations for the combined library approach, we targeted mCardinalΔ19, a near-total tail deletion, with random mutagenesis and isolated six mutations that restored measureable fluorescence and did not hypsochromically shift the emission curve. The six identified error-prone hits together (mCardinal-mut6) restored fluorescence and thermostability nearly back to that of mCardinal. We then built a monomerization library that included the six stabilizing error-prone mutations and a complete tail deletion (Δ20), and that sampled a CPD-generated AC interface and the nine highest-scoring consensus mutations (Table S2). Because the first generation HcRed monomer needed further optimization for improved brightness, we chose to sample a larger surface design landscape than we did in the case of HcRed, again designing the five-residue core of the AC interface, but also allowing diversity in eight other nearby surface positions. The total theoretical library size was 5.7 × 10∧7. After screening 1.1 × 10∧5 variants by fluorescence activated cell sorting (FACS), we isolated two variants that were bright, monomeric, and retained a far-red emission: mKelly1 and mKelly2 (Table 2).

## Discussion

### Clear Design Objectives Speed Protein Development

We demonstrate that an engineering process that makes use of varied protein engineering tools can hasten the isolation of optimized protein variants. Small, focused libraries enriched for diverse but functional HcRed variants addressed separately the problems of brightness, stability, and oligomericity. Beginning with oligomers partially destabilized by the deletion of their C-terminal tails, we quickly moved through functional sequence space, incorporating 38 and 42 mutations over five rounds of design into mGinger1 and mGinger2 respectively (Figures 3 and S4). DNA shuffling (28, 29) enabled us to screen large numbers of candidate mutations in HcRed, but noting that high-value mutations were enriched during this process, we monomerized mCardinal with just one large library, incorporating 39 and 42 mutations respectively into mKelly1 and mKelly2. Importantly, and unlike previous RFP monomerization efforts, we maintained fluorescence at every design stage, allowing us to be stringent in our selections and to maintain far-red emission. The mutations in the final RFP variants were found by employing complementary but divergent engineering processes. Consensus design was used to improve thermo-stability, which has been shown to improve proteins’ evolvability (25, 26), while error-prone mutagenesis added diversity to this pool of stabilizing mutations. Notably, consensus design significantly outperformed random mutagenesis in improving the brightness of HcRed7 (Figure S2). Finally, to build stable and soluble β-sheet surfaces, an application suited neither to consensus design nor error-prone mutagenesis, we used CPD, which we had previously shown to be well suited to this purpose.

**Figure 3.**
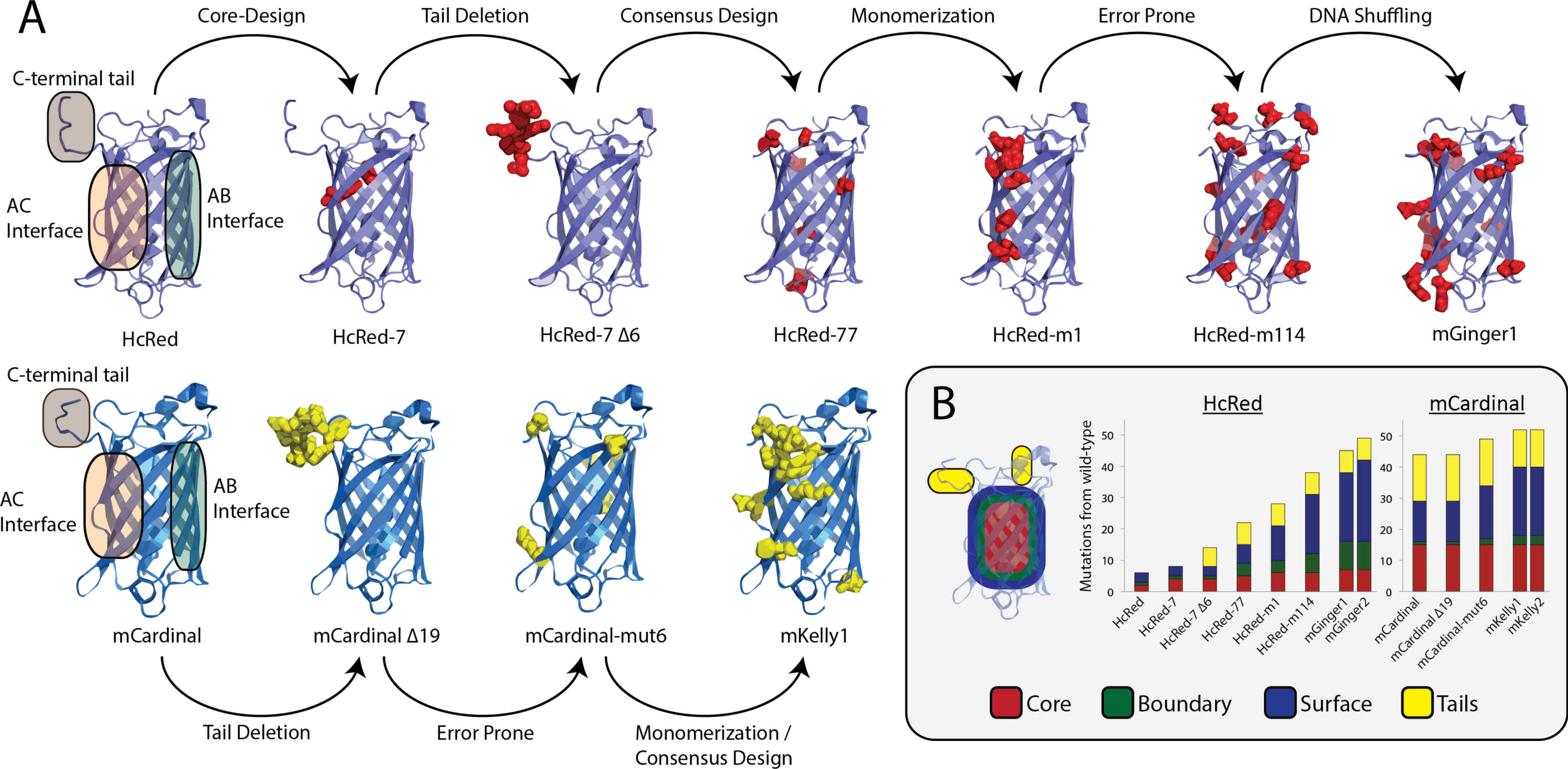
The engineering of the mGingers and mKellys. (A) Each engineering step is shown with the mutations made shown in red spheres in the HcRed background and yellow spheres in the mCardinal background. The crystal structures are of HcRed7 (reported here) and mCardinal (PDB-ID: 4OQW) (B) Solvent-exposed surface area (SASA) was used to categorize the mutations into three buckets, with tails treated separately. The number of mutations separating each designed variant from its wild-type ancestral progenitor (HcCP in the case of HcRed variants and eqFP578 in the case of mCardinal variants) is plotted.

### Mutations Accumulate in Key Structural Regions

A total of 45 mutations in mGinger1 and 52 mutations in mKelly1 separate them from their wild-type progenitors HcCP and eqFP578. These mutations cluster structurally, occurring at the designed AC interface, at chromophore-proximal positions, and near pockets of exposed hydrophobic residues on the protein surface. One region of particular note in which mutations cluster is an apparent channel populated by large numbers of structural water molecules that runs from a wide cleft in the β-barrel between β-strands 7 and 10 to a smaller deformation of β-strands 3 and 11, passing through the chromophore pocket (Figure 2B). These deformations of the β-barrel are bisected by the attachment site of the C-terminal tail, and appear to be stabilized by intermolecular interactions between monomers across the AC interface. A break of the AC interface may destabilize the water channel, putting the chromophore environment into contact with bulk solvent, which would in turn interfere with chromophore maturation and quench fluorescence (23, 30). Indeed, mGinger1 and mKelly1 have eleven and six mutations respectively to residues that are in close proximity (4 Å) to structural waters in this channel and that are not a part of the AC interface (Figure S5). Elsewhere, mGinger1 and mKelly1 have eleven and fifteen mutations respectively to their AC interfaces and two and three to their AB interfaces, breaking oligomerization. In mGinger1 we see eight mutations to patches of exposed hydrophobic surface residues not located at the oligomeric interfaces, as mapped by spatial aggregation propensity (SAP) (31, 32), whereas with mKelly1, we relatively fewer new surface mutations, as we expect that the intense selection and engineering that mCardinal had been subjected to had previously optimized its non-interface surfaces for solubility. Outside of these structural clusters, we introduced relatively few new mutations to mGinger1 and mKelly1, five in each case. mKelly1 does inherit eleven other uncharacterized mutations from mCardinal, to both its surface and core.

### Protein Stability is Linked to Function

Past efforts to monomerize RFPs have ignored the role that scaffold stability may play in engineering a functional monomer. We suggest that as oligomericity is broken, a loss of structural integrity (approximated here by apparent T_m_) can leave single monomers unstable and non-functional. As we monomerized HcRed, we measured the thermal stability of each important intermediate, and found a positive correlation between apparent T_m_ and quantum yield (Figure 4). This relationship may be related to scaffold rigidity, as in a more rigid excited-state chromophore there is less non-radiative decay of fluorescent energy via thermal motion or other atomic interactions. (33, 34). In small molecule fluorophores this is readily seen, as quantum yield increases with decreased temperature (35), and rational design of a chromophore-proximal β-strand was used improve quantum yield of a cyan FP to 0.93 (33). The correlation between quantum yield and apparent T_m_, however, appears to divide into two distinct groups, with dimers having higher quantum yields than monomers. mGinger0.1 and mGinger0.2, for instance, despite being thermostabilized by 5 °C over the parental protein HcRed7, are less bright. We observe an uncoupling between thermostability and quantum yield, where despite sharing an almost identical protein core, the less thermally stable dimer is brighter than its thermostabilized monomeric derivative.

**Figure 4.**
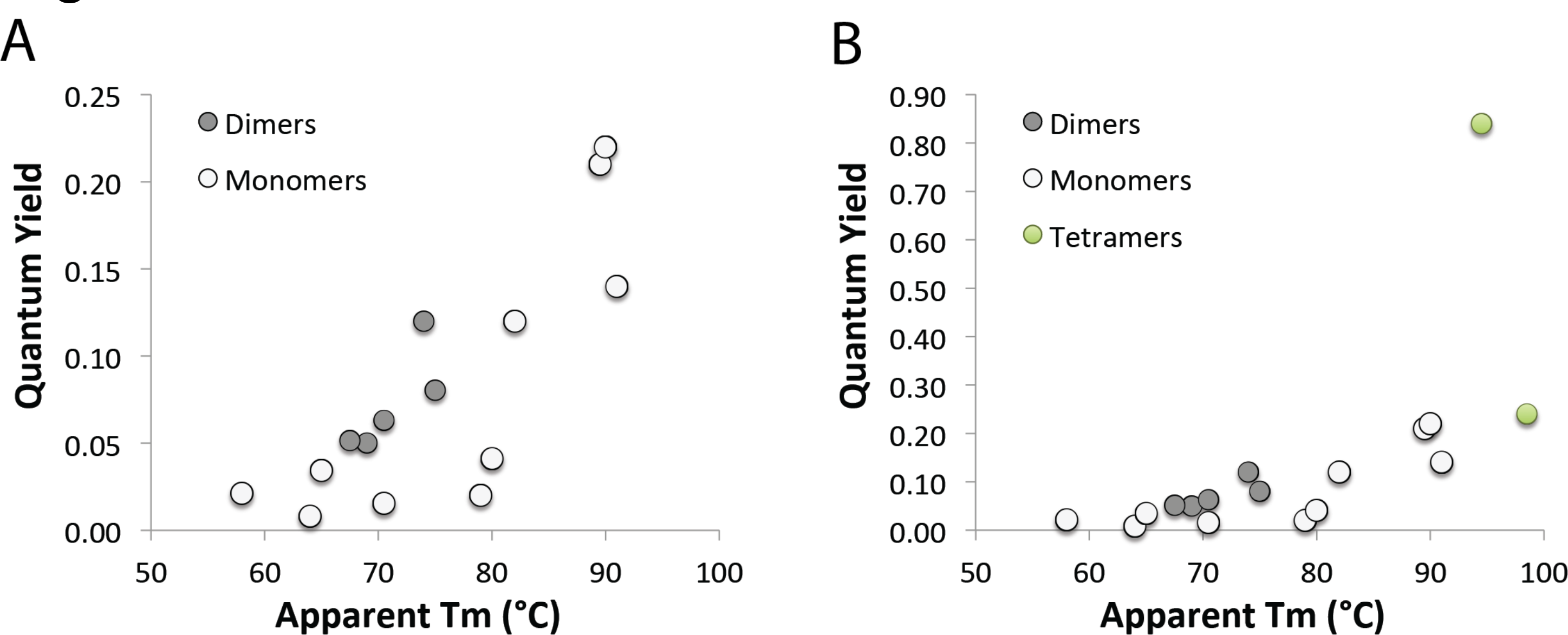
Thermal stability of HcRed, mCardinal, and DsRed variants are plotted against their quantum yields. The correlation can be clearly seen with monomers exhibiting lower quantum yield than dimers at equivalent wavelengths (A) and tetrameric proteins clearly exhibiting the highest quantum yields (B).

### HcRed7’s Structure Explains Brightness and Bathochromic Emission

We solved an x-ray crystal structure of HcRed7, which shows that the mutation from histidine to tyrosine at position 196 serves to add a π-stacking interaction with the chromophore phenolate ring (Figure 1C). Tyr196 π-stacks with the fluorescent cis orientation of the phenolate, serving to both stabilize the fluorescent *cis* phenolate over the non-fluorescent *trans* phenolate—HcRed’s chromophore occupies both *cis* and *trans* conformations—and to red-shift the λ_em_, as a π-stacking phenolate interaction has been shown to reduce the energy of the excited state of the chromophore (36-38). In turn, position 67 is a key catalytic residue that functions as a base, abstracting a proton from the bridging carbon of the phenolate side chain during cyclization (39, 40). This residue is almost invariably a lysine or arginine among RFPs, and we propose that the mutation from arginine to lysine here allows room for the π-stacking interaction and the bulkier tyrosine side chain. It has been previously noted that this π-stacking interaction can induce a bathochromic shift in λ_em_ (41), but here we note that these two mutations also conveyed a 6 °C improvement to apparent Tm and a 60% improvement in quantum yield relative to HcRed.

## Conclusion

We engineered four new monomeric RFPs: mGinger1/2 and mKelly1/2, monomeric variants of the far-red fluorescent proteins HcRed and mCardinal, both dimeric RFPs that had been the targets of unsuccessful monomerization attempts. mKelly1 and mKelly2 join mGarnet and mGarnet2 as part of a new class of bright monomeric RFPs with emission peaking near to or longer than 650 nm (Figure 5). Previously we monomerized DsRed using a pre-stabilized core borrowed from mCherry, and showed that monomerization is possible with little to no change to an RFP’s spectroscopic properties (19). Here we show that stabilization of the entire protein scaffold is important for monomerization. Despite the mGingers and mKellys being slightly dimmer and hypsochromically shifted from HcRed7 and mCardinal, they move the needle toward longer wavelengths and brighter emission. Past monomerization efforts have been beset by similar loss of brightness, but because they necessitated significant mutation to the core of the protein and the chromophore environment (13, 15, 42-44), it has been difficult to separate the effects of potentially suboptimal core mutation from the inescapable externalities of monomerization. The rational approach that we lay out in monomerizing HcRed and mCardinal includes elements of rational design, computational design, and directed evolution, and represents a marked improvement in both the speed and efficiency of RFP monomerization. Further exploration of stable RFP cores will be necessary to determine how to significantly improve brightness post-monomerization.

**Figure 5.**
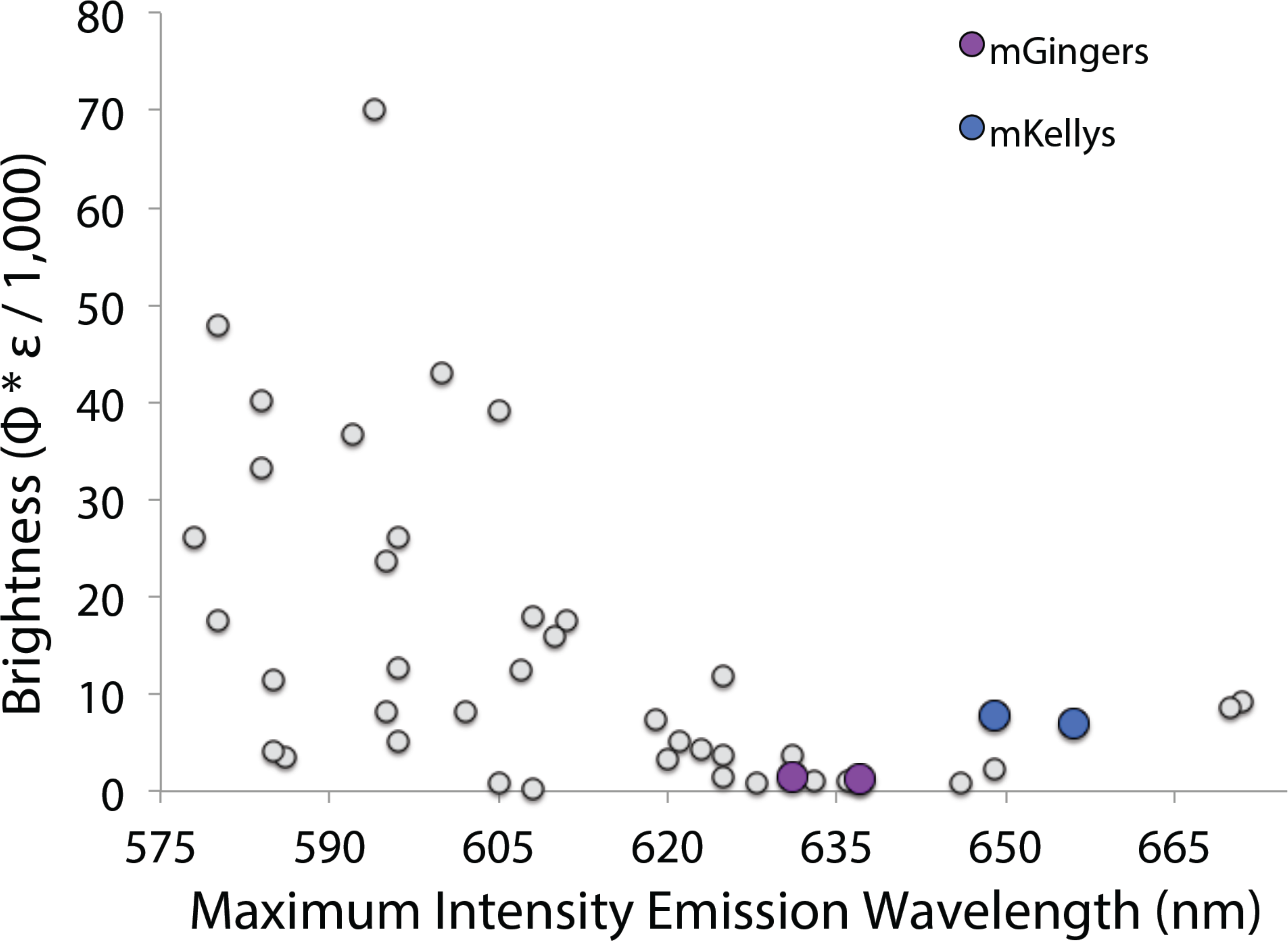
There is a negative correlation between brightness and λ_em_ among RFPs. All known monomeric RFPs whose brightness and λ_em_ have been measured are plotted. The mGingers and mKellys are among the furthest red-shifted monomeric proteins described to date.

## Materials and Methods

### Plasmids and Bacterial Strains

The HcRed sequence was taken and modified from the HcCP Genbank entry (accession number AF363776). Ten amino acids were added to the N-terminus, consisting of a Methionine followed by a 6x Histidine tag for protein purification, followed by a Gly-Ser-Gly linker sequence. All genes were constructed by overlap extension PCR from oligonucleotides designed by DNAworks and ordered from Integrated DNA Technologies (IDT). Assembled genes were PCR-amplified and cloned into the pET-53-DEST expression plasmid (EMD Millipore). Constructs were sequence-verified and transformed into BL21-Gold(DE3) competent cells for protein expression (Agilent).

### Construction of Designed Libraries

The HcRed core variants were designed with DNAworks as “mutant runs” of the wild-type gene assembly. The HcRed AC surface library used the triplet codon “VRN” to replace the five design positions, which allows for the possible amino acids D/E/G/H/K/N/Q/R/S. The mCardinal monomer library was designed by hand, as it was too complex for DNAworks; we used degenerate bases where possible. For all libraries, oligonucleotides were ordered from IDT and cloning was carried out as described above.

### Error Prone Mutagenesis

Error prone mutagenesis of HcRed variants was performed by addition of manganese chloride to Taq DNA polymerase PCR reactions. 10µM, 15µM, and 20µM MnCl_2_ were tested and cloned with PIPE cloning into pET-53-DEST for sequencing. Twelve colonies from each library were picked and sequenced, and the library with a mutation rate closest to but not more than 2.0 mutations per gene was selected for further screening.

### DNA Shuffling

The variants that were to be shuffled together were PCR-amplified and purified by gel electrophoresis with a standard spin-column gel purification kit (Qiagen). 5 µg of the purified DNA was then digested with 0.5 U of DNAseI (NEB) in a 50 µl reaction. The reaction was allowed to sit for 7.5 minutes at room temperature and then quenched with 5 µl of 100 mM EDTA (4x the concentration of MgCl_2_ in the reaction buffer). The reaction was further heat-inactivated for 10 minutes at 90ºC in a thermocycler and electrophoresed. Bands of ~30 bp, as compared to standards [30 bp oligo (IDT) / 100 bp DNA ladder (NEB)], were excised, frozen, and then purified using a Freeze ‘N Squeeze gel purification kit (BioRad) because the small band size precluded spin column purification. Purified digested fragments were mixed together at a 1:1 ratio and assembled via overlap-extension PCR.

### Generating a multiple sequence alignment (MSA) and computing a consensus sequence

We searched various resources including GenBank, SwisProt, UniProt, NCBI-BLAST, and patent databases for reported FP sequences. We found 741 unique fluorescent protein sequences and aligned them with MAFFT, which we then hand-curated with the use of a 163-member structural alignment. Each position in the resulting alignment was scored as follows (45, 46). First, sequences were Henikoff weighted (47) to account for the presence of highly similar sequences in the MSA. The entropy *H_C_
* of each column in the alignment was then calculated, ignoring any gaps. Finally, a score was obtained for each non-gap character in each column via:

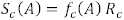

Here, *f_C_
*(*A*) is the frequency of character *A* in column *C*, and *R_C_
* is the uncertainty reduction in each column (corrected for the column gap fraction *f_C_
* (–)) calculated as:

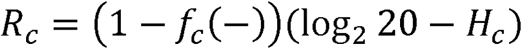

The highest scoring character in each column (or multiple characters in the case of ties) was selected as the consensus amino acid at that position.

### Protein Expression and Library Screening

Single bacterial colonies were picked with sterile toothpicks and inoculated into 300 µl of Super Optimal Broth (SOB) supplemented with 100 µg/ml ampicillin in 2 ml deep-well 96-well plates (Seahorse Biosciences). The plates were sealed with microporous film (Denville Scientific) to facilitate gas exchange during growth. Cultures were grown overnight at 37 °C / 300 RPM. The next morning 800 µl of fresh SOB with 100 µg/ml ampicillin and 1mM Isopropyl β-D-1-thiogalactopyranoside (IPTG) was added to a total volume of 1 ml (evaporation losses overnight are approximately 100 µl). Plates were then shaken 12 hours at either 30ºC or 37 °C and 400 RPM. After overnight expression, plates were screened with a liquid handling robot (Tecan Freedom Evo) linked to a platereader (Tecan Saffire 2). 200 µl of each culture was added to Greiner UV-Star 96-well plates and imaged for fluorescence emission at 675 nm after excitation at 600 nm. Controls were included on each plate to account for plate-to-plate variation. Potential hits were streaked out onto a fresh LB-Amp plate, grown overnight at 37ºC, and four colonies were picked for each potential hit. These were then grown again and screened as detailed above, with hits then ranked on their significant variation from the parent or control.

### Protein Purification

To further characterize important variants, 1 L of SOB in Fernbach flasks was inoculated 1:100 with overnight cultures, grown to an OD of ~0.5 and induced at 37ºC for 12 hours with 1mM IPTG. The broth was then transferred to centrifuge flasks and spun at 5,000 × g in a fixed angle rotor for 10 min and the supernatant decanted. Bacterial pellets were resuspended in 25 ml of lysis buffer (50 mM sodium phosphate, 150 mM NaCl, 0.1% v/v Triton-X, pH 7.4) supplemented with 50 Units/ml Benzonase (Sigma) and 0.05 mg/ml Hen Egg Lysozyme (Sigma). Resuspended pellets were then run over a microfluidizer to fully lyse the bacteria. To pellet down the cellular debris, the lysed cultures were again centrifuged for 10 minutes at 15,000 × g in a fixed angle rotor. The colored supernatant was then poured through a column of His-Select resin (Sigma), washed twice (50 mM sodium phosphate, 150 mM NaCl, 15 mM Imidazole, pH 7.4), and eluted with 500 µl elution buffer (50 mM sodium phosphate, 150 mM NaCl, 250 mM Imidazole, pH 7.4). Proteins were further purified by FPLC (AKTA) with a Superdex 75 10/300 column, and in the process buffer exchanged into PBS.

### Fluorescent Protein Characterization

Purified protein variants were assayed in triplicate in Greiner UV-Star 96-well plates with a Tecan Saffire 2. An absorbance scan (260 – 650 nm), a fluorescence excitation scan (500 – 640 nm excitation / 675 nm emission), and a fluorescence emission scan (550 nm excitation / 575 – 800 nm emission) were run on 100 µl of eluted protein to determine spectral peaks.

To measure the quantum yield we diluted each protein so that the absorbance for 200 µl of protein at 540 nm was between 0.1 and 0.5. We then measured the A_550_ in triplicate (or duplicate if it was a poorly expressed protein), diluted the sample to an A_550_ of 0.04 and took an emission scan (540 nm excitation / 550 – 800 nm emission). The area under the emission curve was calculated after fitting it to a 4^th^ order Gaussian, and the quantum yield was calculated with the following formula:

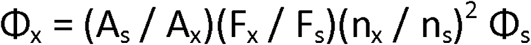

Where Φ is quantum yield, A is absorbance, F is total fluorescent emission (area under the curve), and n is the refractive index of the solvents used. Subscript X refers to the queried substance and subscript S refers to a standard of known quantum yield. It is important that the standard be excited with the same wavelength of light as the unknown sample. We use DsRed, which has a known quantum yield of 0.79 as the protein standard.

To measure extinction coefficient we took 100 µl of the protein solution that had been diluted to an A_550_ of between 0.1 and 0.5 and measured absorbance between 400 nm and 700 nm in triplicate. We then added 100 µl of 2M NaOH to each well and remeasured absorbance between 400 nm and 700 nm. The base-denatured chromophore, which peaks at approximately 450 nm has a known extinction coefficient of 44,000 M^-1^cm^-1^. Then to calculate the extinction coefficient is calculated with the following formula:

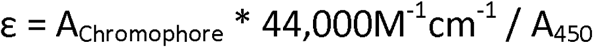

### Thermal Stability

Purified proteins were diluted to an absorbance of 0.2 at the wavelength of maximum absorbance (λ_abs_) so that their fluorescence would not saturate the rtPCR detector. 50 µl of each purified protein was then loaded into a 96-well PCR plate and covered with clear optical tape. The proteins were incubated at 37°C for 10 minutes and then the temperature was ramped at 0.5°C every 30 seconds up to 99°C, with fluorescence measured every ramp step in a CFX96 Touch Real-Time PCR Detection System (Bio-Rad). We refer to this as a thermal melt. The derivative curve of the thermal melt finds the inflection point of the slope, which is the apparent temperature at which fluorescence is irrevocably lost (apparent Tm).

### Oligomeric Determination

(A) Size exclusion chromatography. 100 µl of each purified protein analyzed was run over a Superdex 75 10/300 size exclusion column with 25 ml bed volume on an AKTA from GE Life Sciences. Absorbance was measured after passage through the column at 575 nm, where the red chromophore absorbs. (B) Analytical ultracentrifugation. Purified protein samples were diluted to an A_575_ of 0.5 for a path-length of 1.25 cm. These samples were put into two-channel sedimentation velocity cuvettes with the blank channel containing PBS. Sedimentation velocity was run at 40,000 RPM overnight with full A_575_ scans collected with no pause between reads. Data was loaded into Sedfit and a c(m) distribution was run with default assumptions made for PBS buffer viscosity. After integration, the c(m) curve was exported to Excel. (C) Homo-FRET. 200 µl of each purified protein was diluted to an Absorbance of 0.1 to 0.5 at 530 nm in 96-well Greiner UV-Star plates. Polarization scans were then taken with excitation at 530 nm and emission at 610 nm in a Tecan Safire2 plate-reader. Rose Bengal was used as a standard to calculate the instrument G factor (mP = 349).

### Crystallography

Rectangular plate crystals of HcRed7 grew in 7 days by the sitting-drop vapor diffusion method in 100 mM Bis-Tris pH 6.5 with 200 mM ammonium sulfate and 25% w/v PEG 3350. Crystals were flash frozen in 2-Methyl-2,4-pentanediol (MPD) and shipped to beamline 12-2 at the Stanford Synchrotron Radiation Lightsource, where a 1.63 Å data set was collected. Phases were obtained through molecular replacement using the crystal structure of HcRed (PDB ID 1YZW).

Following molecular replacement, model building and refinement were run with COOT and PHENIX.(48, 49) NCS restraints were applied to early refinement steps and removed at the final stages of refinement. TLS parameters were used throughout. The chromophore was initially left out of the refinement and added at a later stage when clear density became evident for it. Coordinates were deposited in the Protein Data Bank with the code [to be submitted]. Data collection and refinement statistics are listed in Table S1.

## Acknowledgments

Helpful conversations were had with Yun Mou, Matthew Moore, and Roberto Chica. We would also like to thank Professor Frances Arnold for her advice and support. Finally, we would like to acknowledge the funding and support of the National Institute of Biomedical Imaging and Bioengineering (NIBIB).

---

**Figure S1.**
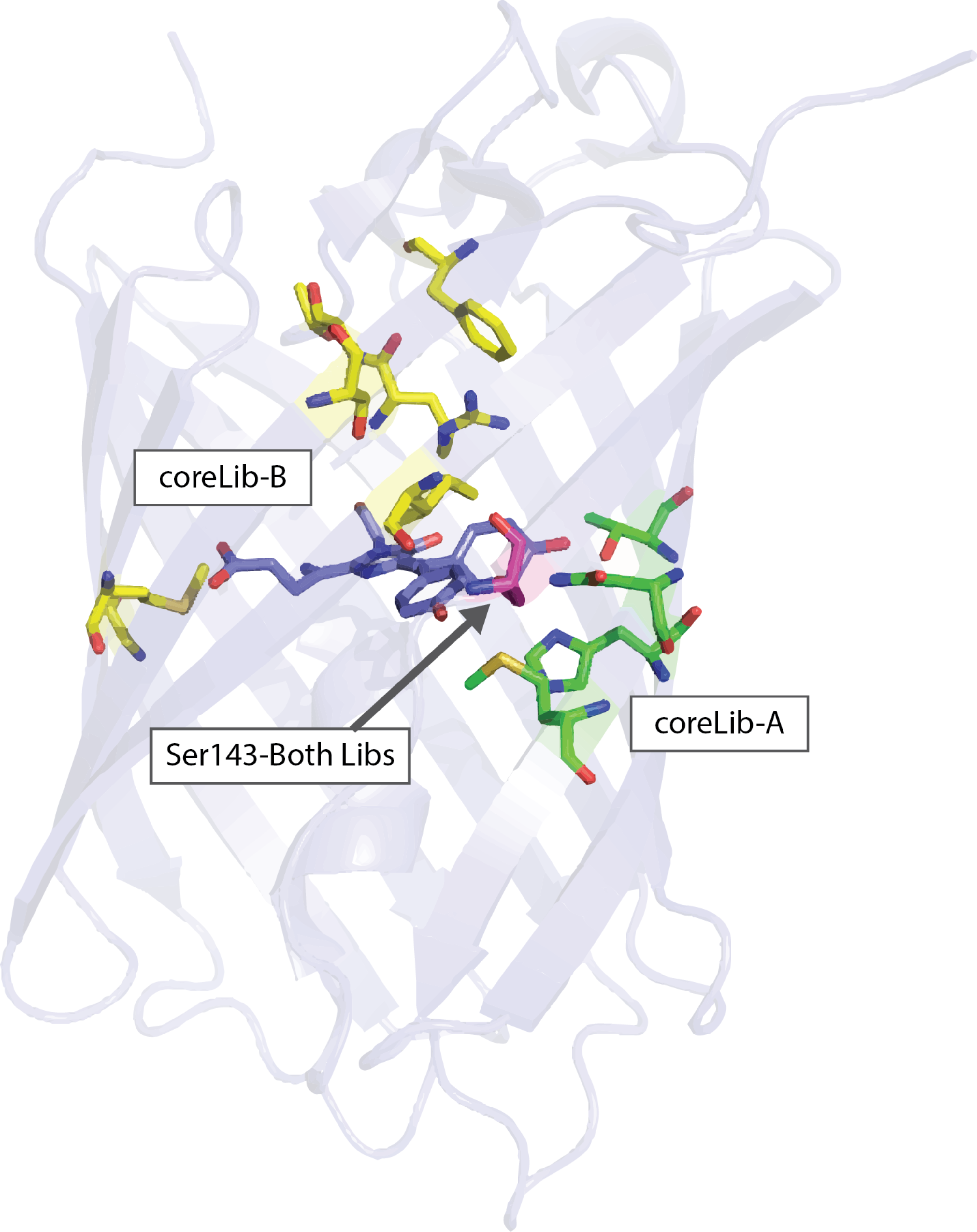

**Figure S2.**
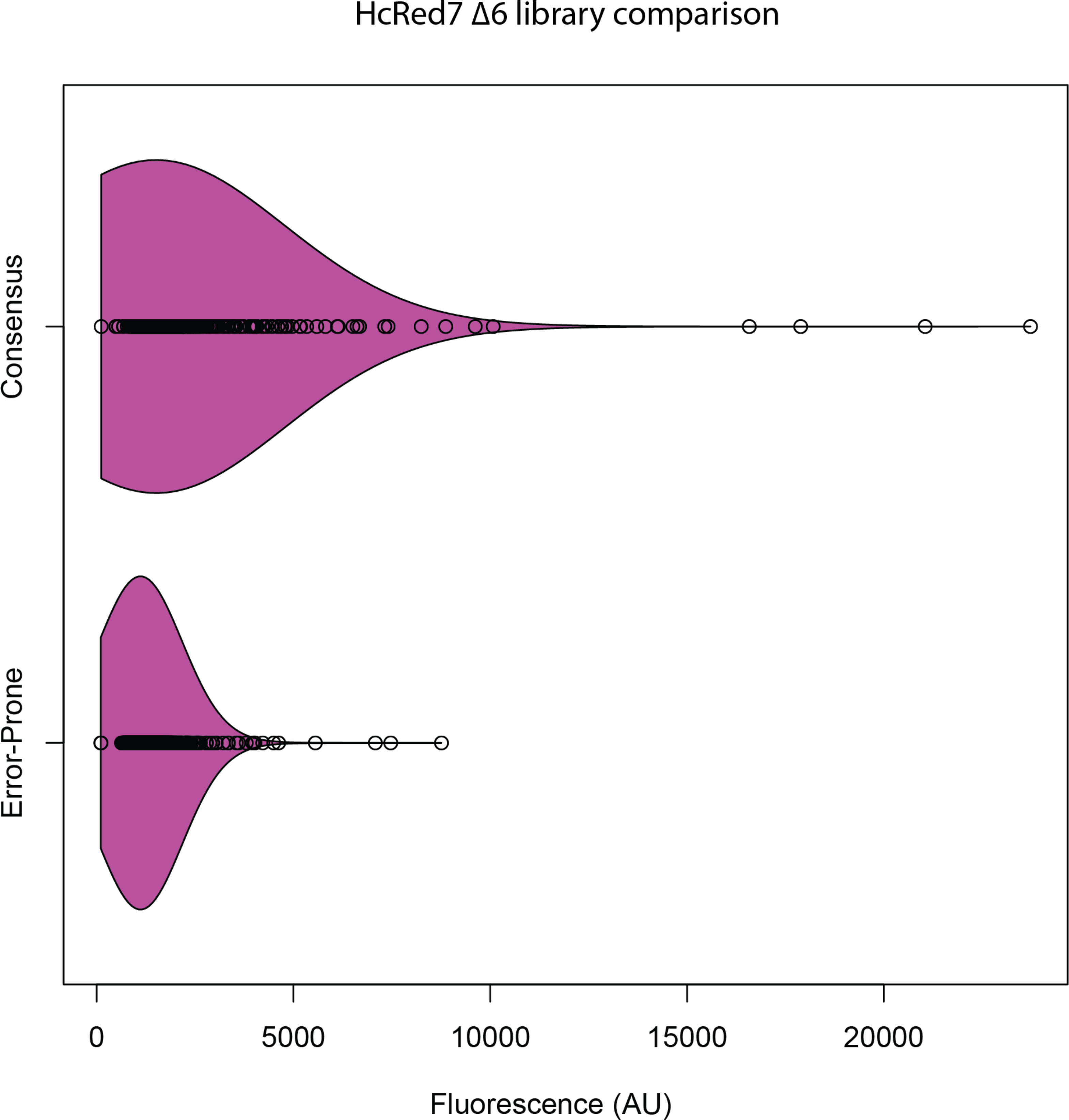

**Figure S3.**
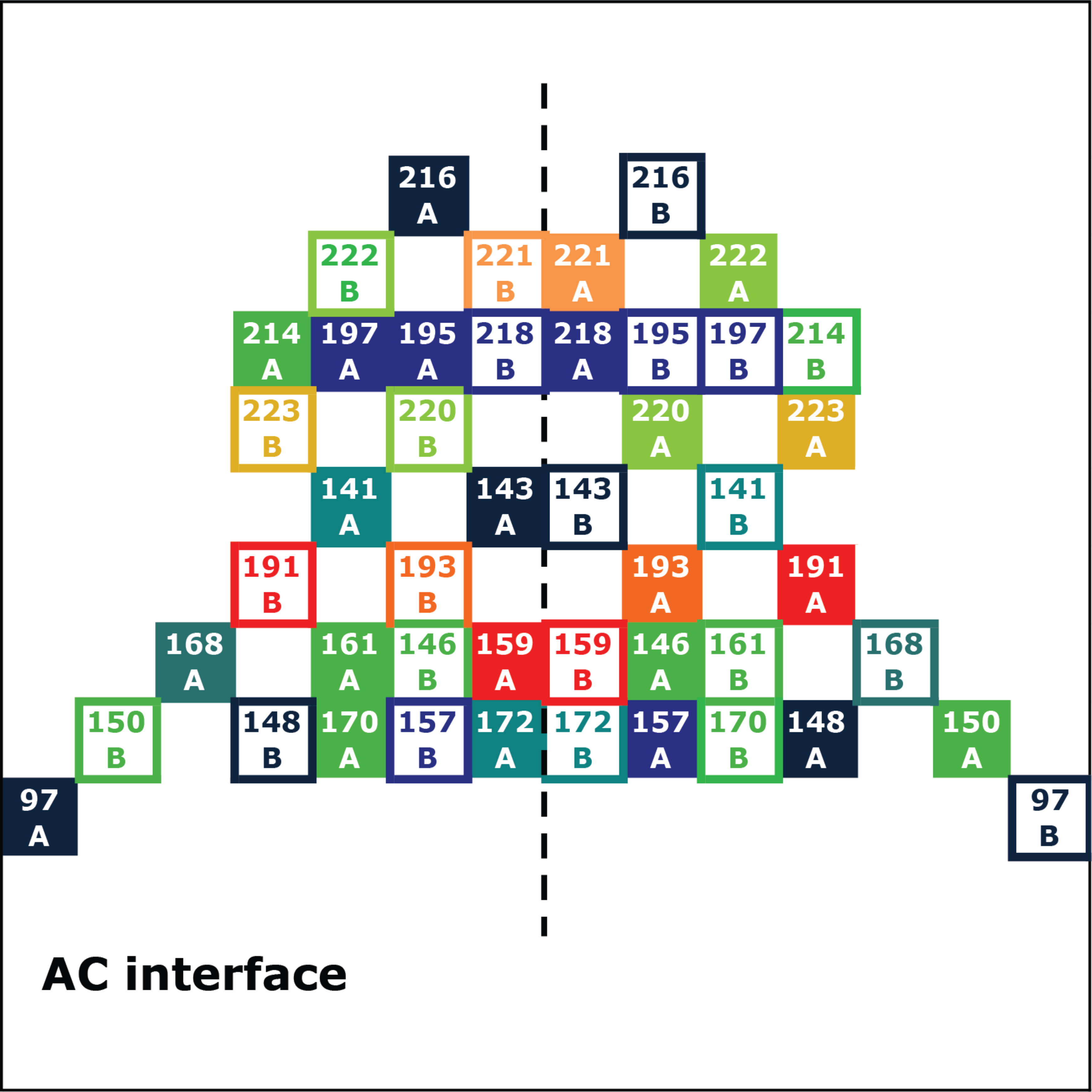

**Figure S4.**
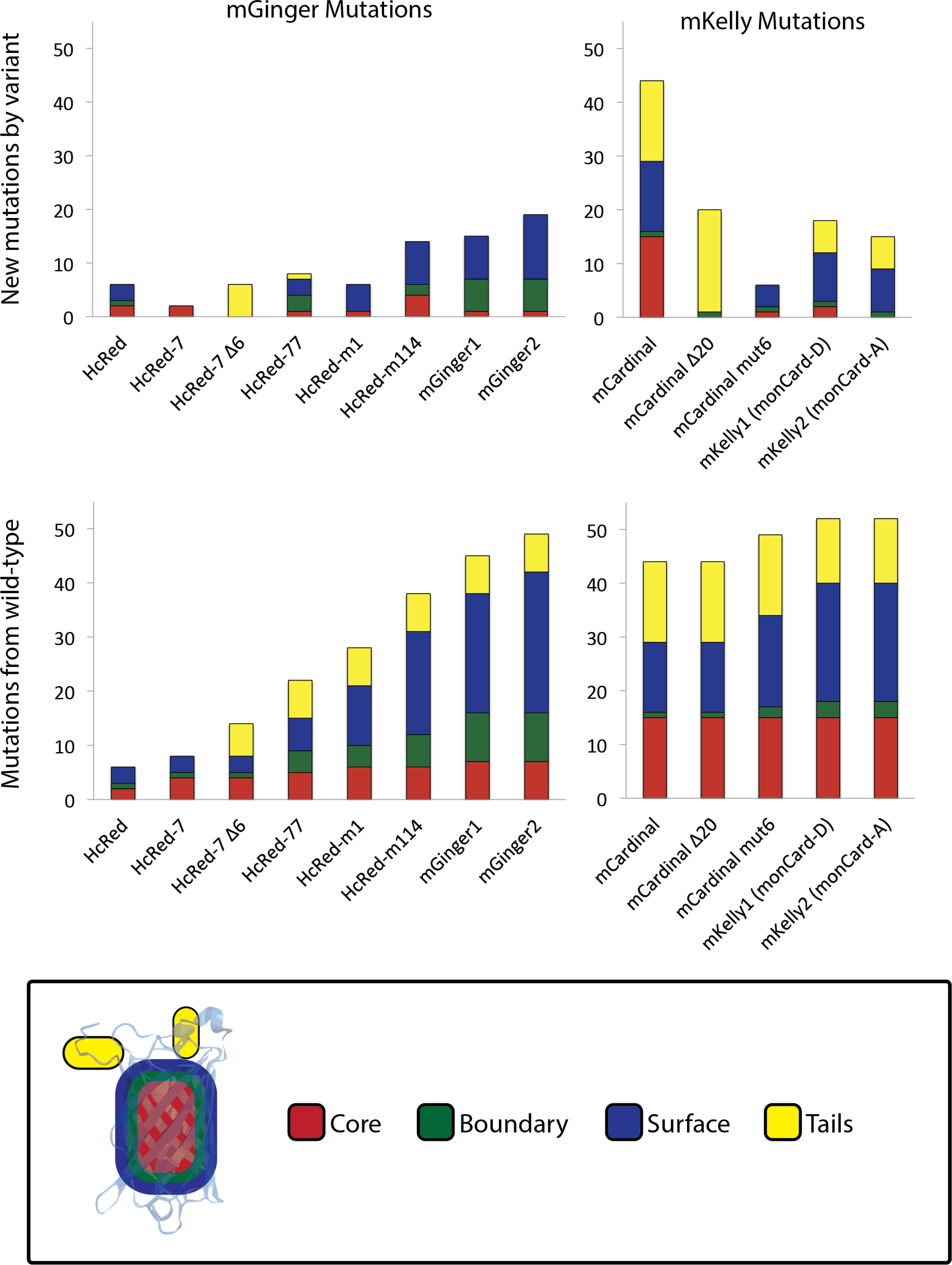

**Figure S5.**
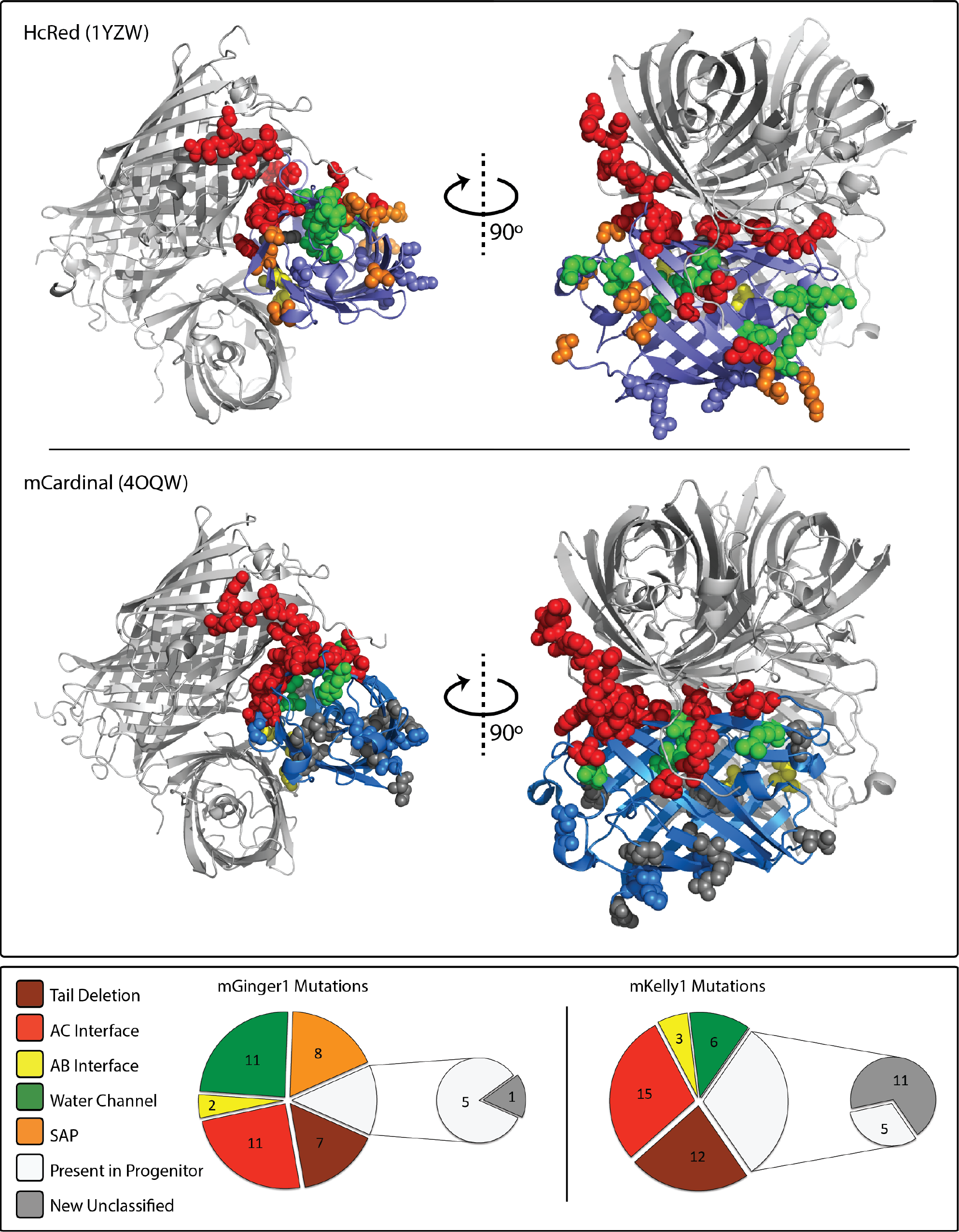

## References

1. Matela G , Gao P , Guigas G , & Eckert AF (2017) A far-red emitting fluorescent marker protein, mGarnet2, for microscopy and STED nanoscopy. Chemical ….

2. Tromberg BJ , et al. (1999) Non-invasive in vivo characterization of breast tumors using photon migration spectroscopy. Neoplasia (New York, N.Y.) 2(1-2):26–40.

3. Bashkatov AN , Genina EA , Kochubey VI , & Tuchin VV (2005) Optical properties of human skin, subcutaneous and mucous tissues in the wavelength range from 400 to 2000 nm. Journal of Physics D: Applied Physics 38(15):2543.

4. Wang S , Moffitt JR , Dempsey GT , Xie XS , & Zhuang X (2014) Characterization and development of photoactivatable fluorescent proteins for single-molecule–based superresolution imaging. Proceedings of the National Academy of Sciences of the United States of America.

5. Wenqin W , Gene-Wei L , Chongyi C , Xie XS , & Xiaowei Z (2011) Chromosome Organization by a Nucleoid-Associated Protein in Live Bacteria. Science 333(6048):1445–1449.

6. Filonov GS , et al. (2011) Bright and stable near-infrared fluorescent protein for in vivo imaging. Nature biotechnology 29(8):757–761.

7. Alieva NO , et al. (2007) Diversity and evolution of coral fluorescent proteins. PloS ONE 3(7).

8. Shagin DA , et al. (2004) GFP-like proteins as ubiquitous metazoan superfamily: evolution of functional features and structural complexity. Molecular Biology and Evolution 21(5):841–850.

9. Bou-Abdallah F , Chasteen ND , & Lesser MP (2006) Quenching of superoxide radicals by green fluorescent protein. Biochimica et Biophysica Acta (….

10. Palmer CV , Modi CK , & Mydlarz LD (2009) Coral Fluorescent Proteins as Antioxidants. PLoS ONE 4(10).

11. Salih A , Larkum A , Cox G , Kühl M , & Hoegh-Guldberg O (2000) Fluorescent pigments in corals are photoprotective. Nature 408(6814):850–853.

12. Smith EG , D’Angelo C , Salih A , & Wiedenmann J (2013) Screening by coral green fluorescent protein (GFP)-like chromoproteins supports a role in photoprotection of zooxanthellae. Coral reefs.

13. Campbell RE , et al. (2002) A monomeric red fluorescent protein. Proceedings of the National Academy of Sciences.

14. Kredel S , et al. (2008) mRuby, a bright monomeric red fluorescent protein for labeling of subcellular structures. PloS ONE 4(2).

15. Shemiakina II , et al. (2012) A monomeric red fluorescent protein with low cytotoxicity. Nature Communications.

16. Takako K , et al. (2006) A fluorescent variant of a protein from the stony coral Montipora facilitates dual-color single-laser fluorescence cross-correlation spectroscopy. Nature Biotechnology 24(5):577–581.

17. Gurskaya NG , et al. (2001) GFP-like chromoproteins as a source of far-red fluorescent proteins1. FEBS Letters 507(1):16–20.

18. Subach OM , et al. (2008) Conversion of Red Fluorescent Protein into a Bright Blue Probe. Chemistry & Biology 15(10):1116–1124.

19. Wannier TM , Moore MM , Mou Y , & Mayo SL (2015) Computational Design of the β-Sheet Surface of a Red Fluorescent Protein Allows Control of Protein Oligomerization. PloS one 10(6).

20. Wall MA , Socolich M , & Ranganathan R (2000) The structural basis for red fluorescence in the tetrameric GFP homolog DsRed. Nature Structural Biology 7(12):1133–1138.

21. Wannier TM & Mayo SL (2014) The structure of a far-red fluorescent protein, AQ143, shows evidence in support of reported red-shifting chromophore interactions. Protein Science 23(8):11481153.

22. Mudalige K , et al. (2010) Photophysics of the red chromophore of HcRed: evidence for cis-trans isomerization and protonation-state changes. The journal of physical chemistry. B 114(13):4678–4685.

23. Moore MM , Oteng-Pabi SK , Pandelieva AT , Mayo SL , & Chica RA (2012) Recovery of Red Fluorescent Protein Chromophore Maturation Deficiency through Rational Design. PLoS ONE 7(12).

24. Piatkevich KD , et al. (2012) Extended Stokes shift in fluorescent proteins: chromophore-protein interactions in a near-infrared TagRFP675 variant. Scientific reports 3:1847.

25. Bloom JD , Labthavikul ST , Otey CR , & Arnold FH (2006) Protein stability promotes evolvability. Proceedings of the National Academy of Sciences of the United States of America 103(15):5869–5874.

26. Jäckel C , Bloom JD , Kast P , Arnold FH , & Hilvert D (2010) Consensus protein design without phylogenetic bias. Journal of molecular biology 399(4):541–546.

27. Pletneva NV , et al. (2011) Crystallographic study of red fluorescent protein eqFP578 and its far - red variant Katushka reveals opposite pH - induced isomerization of chromophore. Protein Science 20(7):1265–1274.

28. Crameri A , Whitehorn EA , Tate E , & Stemmer WP (1996) Improved green fluorescent protein by molecular evolution using DNA shuffling. Nature biotechnology 14(3):315–319.

29. Zhao H & Arnold FH (1997) Optimization of DNA Shuffling for High Fidelity Recombination. Nucleic Acids Research.

30. Regmi C, K. , Bhandari YR , Gerstman BS , & Chapagain, PP . (2013) Exploring the Diffusion of Molecular Oxygen in the Red Fluorescent Protein mCherry Using Explicit Oxygen Molecular Dynamics Simulations. The journal of physical chemistry. B.

31. Black SD & Mould DR (1991) Development of hydrophobicity parameters to analyze proteins which bear post- or cotranslational modifications. Analytical Biochemistry 193(1):72–82.

32. Chennamsetty N , Voynov V , Kayser V , Helk B , & Trout BL (2010) Prediction of aggregation prone regions of therapeutic proteins. The journal of physical chemistry. B 114(19):6614–6624.

33. Goedhart J , et al. (2011) Structure-guided evolution of cyan fluorescent proteins towards a quantum yield of 93%. Nature communications 3:751.

34. Lelimousin M , et al. (2009) Intrinsic dynamics in ECFP and Cerulean control fluorescence quantum yield. Biochemistry 48(42):10038–10046.

35. Rurack K & Spieles M (2011) Fluorescence quantum yields of a series of red and near-infrared dyes emitting at 600-1000 nm. Analytical chemistry 83(4):1232–1242.

36. Chica RA , Moore MM , Allen BD , & Mayo SL (2010) Generation of longer emission wavelength red fluorescent proteins using computationally designed libraries. Proceedings of the National Academy of Sciences of the United States of America 107(47):20257–20262.

37. Morozova KS , et al. (2010) Far-red fluorescent protein excitable with red lasers for flow cytometry and superresolution STED nanoscopy. Biophysical journal 99(2):5.

38. Grigorenko BL , Nemukhin AV , Polyakov IV , & Krylov AI (2013) Triple-decker motif for red-shifted fluorescent protein mutants. The Journal of Physical Chemistry Letters 4(10):1743–1747.

39. Strack RL , Strongin DE , Mets L , Glick BS , & Keenan RJ (2010) Chromophore formation in DsRed occurs by a branched pathway. Journal of the American Chemical Society 132(24):8496–8505.

40. Tubbs JL , Tainer JA , & Getzoff ED (2005) Crystallographic Structures ofDiscosomaRed Fluorescent Protein with Immature and Mature Chromophores: Linking Peptide BondTrans−CisIsomerization and Acylimine Formation in Chromophore Maturation†,‡. Biochemistry 44(29):9833–9840.

41. Pandelieva AT , et al. (2015) Brighter Red Fluorescent Proteins by Rational Design of Triple-Decker Motif. ACS chemical biology.

42. Strongin DE , et al. (2007) Structural rearrangements near the chromophore influence the maturation speed and brightness of DsRed variants. Protein Engineering, Design & Selection 20(11):525–534.

43. Henderson JN , et al. (2009) Excited State Proton Transfer in the Red Fluorescent Protein mKeima. Journal of the American Chemical Society 131(37):13212–13213.

44. Kredel S , et al. (2009) mRuby, a Bright Monomeric Red Fluorescent Protein for Labeling of Subcellular Structures. PLoS ONE 4(2).

45. Schneider TD (2002) Consensus sequence zen. Applied bioinformatics.

46. Schneider TD & Stephens RM (1990) Sequence logos: a new way to display consensus sequences. Nucleic acids research.

47. Henikoff S & Henikoff JG (1994) Position-based sequence weights. Journal of molecular biology.

48. Adams P , et al. (2010) PHENIX: a comprehensive Python-based system for macromolecular structure solution. Acta crystallographica. Section D, Biological crystallography 66(Pt 2):213–221.

49. Emsley P , Lohkamp B , Scott W , & Cowtan K (2010) Features and development of Coot. Acta crystallographica. Section D, Biological crystallography 66(Pt 4):486–501.

